# Identifying individual risk rare variants using protein structure-guided local tests (POINT)

**DOI:** 10.1101/333245

**Authors:** Rachel Marceau West, Wenbin Lu, Daniel M. Rotroff, Melaine Kuenemann, Sheng-Mao Chang, Michael J. Wagner, John B. Buse, Alison Motsinger-Reif, Denis Fourches, Jung-Ying Tzeng

## Abstract

Rare variants are of increasing interest to genetic association studies because of their etiological contributions to human complex diseases. Due to the rarity of the mutant events, rare variants are routinely analyzed on an aggregate level. While aggregation analyses improve the detection of global-level signal, they are not able to pinpoint causal variants within a variant set. To perform inference on a localized level, additional information, e.g., biological annotation, is often needed to boost the information content of a rare variant. Following the observation that important variants are likely to cluster together on functional domains, we propose a <u>p</u>r<u>o</u>tei<u>n</u> structure guided local <u>t</u>est (POINT) to provide variant-specific association information using structure-guided aggregation of signal. Constructed under a kernel machine framework, POINT performs local association testing by borrowing information from neighboring variants in the 3-dimensional protein space in a data-adaptive fashion. Besides merely providing a list of promising variants, POINT assigns each variant a p-value to permit variant ranking and prioritization. We assess the selection performance of POINT using simulations and illustrate how it can be used to prioritize individual rare variants in *PCSK9* associated with low-density lipoprotein in the Action to Control Cardiovascular Risk in Diabetes (ACCORD) clinical trial data.

**Author summary:** While it is known that rare variants play an important role in understanding associations between genotype and complex diseases, pinpointing individual rare variants likely to be responsible for association is still a daunting task. Due to their low frequency in the population and reduced signal, localizing causal rare variants often requires additional information, such as type of DNA change or location of variant along the sequence, to be incorporated in a biologically meaningful fashion that does not overpower the genotype data. In this paper, we use the observation that important variants tend to cluster together on functional domains to propose a new approach for prioritizing rare variants: the <u>p</u>r<u>o</u>tei<u>n</u> structure guided local <u>t</u>est (POINT). POINT uses a gene’s 3-dimensional protein folding structure to guide aggregation of information from neighboring variants in the protein in a robust manner. We show how POINT improves selection performance over single variant tests and sliding window approaches. We further illustrate how it can be used to prioritize individual rare variants using the Action to Control Cardiovascular Risk in Diabetes (ACCORD) clinical trial data, finding five promising variants within *PCSK9* in association with low-density lipoprotein, including three new mutations near the PCSK9-LDLR binding domain.

## Introduction

Rare genetic variants, e.g. those which occur in less than 1-3% of a population, play an important role in complex diseases. Individual rare variants can be difficult to detect due to low frequencies of the mutant alleles. Therefore, associations involving rare variants are typically discerned using “global” or variant-set tests, which aggregate information across variants to gain sufficient power. These aggregation tests can be done in a burden-based fashion (i.e., modeling phenotype as a function of a weighted sum of genetic markers) [1–4], or using kernel tests (i.e., examining association between pairwise trait similarity and pairwise genetic similarity) [5–9]. Global aggregation tests substantially improve the power for detecting set-level association with phenotypes; however, they are not able to identify individual rare risk variants responsible for the set-level significance.

Localizing rare risk variants from a significant variant set can help guide follow-up studies and provide insight into the functionality and molecular mechanisms of the phenotypes. Several methods have been proposed to prioritize individual rare risk variants based on single-variant analysis [10–12]; yet it has been shown that borrowing external information, either from biological annotations or from other rare variants, can amplify the information content, better separate causal and non-causal variants, and significantly stabilize inferences made at the local level [13].

One approach for variant prioritization involves using functional annotation to filter out variants that are less likely to be causal [14, 15]. Informative functional annotation may include variant frequency, type of DNA change (e.g., frameshift, missense, etc.), conservation score, and predicted impact of the variant on protein structure and gene constraint [15]. While useful for providing a subset of likely causal variants, annotation-based filtering is often phenotype non-specific, and may lead to high false negative selection rates when rigid variant-exclusion thresholds are applied based on one or more filtering criteria [15].

A second class of prioritization methods incorporates functional information as a prior to avoid using absolute rules to include or exclude variants. These functional priors, typically imposed on variant effects, have been included in hierarchical modeling frameworks [13, 16, 17] and Bayesian variable selection models [18, 19]. Methods of this type reduce the occurrence of false negatives as described above and allow the trait-variant association to guide variant selection, yielding better prioritization performance. In addition, these hierarchical approaches facilitate estimation of individual effects of the rare variants. However, these methods can be computationally demanding as the computational burden grows with increasing numbers of variants.

A third class of prioritization methods searches for genomic clustering of rare risk variants. These methods stem from the observation that functional or disease-causing variants are more likely to cluster together than null variants [20–23] in the functional domains. Yue et al. [20] note the existence of “domain hotspots”, or mutational hotspots, within which known functionally significant mutations are more likely to cluster together compared to random nonsynonymous variants. Frank et al. [21] discuss significant clusters of variants within glutamate domains in schizophrenia and bipolar disorder. It has also been shown that actions and interactions of regulatory elements (e.g., promoters, repressors, and enhancers) may be one key reason for relevant loci to cluster within functional domain or mutational hotspots [22]. Based on the observance of domain hotspots, various methods have been proposed to exhaustively search for the single nucleotide polymorphism (SNP) subset that is most significantly associated with the phenotype, either in 2-dimensional (2D) sequence space [24–27] or among all possible SNP subsets [28–31]. All-subset searches may provide better coverage, especially when risk variants do not cluster closely together in the 2D sequence space (such as in the case of regulatory elements). However, the computational burden of an all-subset search can be intractable when a large number of variants are of interest, and consequently require splitting up the target genomic region into segments beforehand [29], which may lead to missing an optimal subset split over arbitrarily defined segments.

In this work, we propose the **<u>p</u>**r**<u>o</u>**te**<u>in</u>** structure guided local **<u>t</u>**est (POINT) as a new method for prioritizing individual risk rare variants. Like the third class of prioritization methods which focuses on genomic clusters to pinpoint rare causal variants, POINT is built upon the rationale that risk variants tend to cluster within functional domains or mutational hotspots [20–23]. In order to search beyond the 2D sequence space yet retain computational efficiency, however, POINT relies on the tertiary protein structure, i.e., the 3-dimensional (3D) folding of amino acids, to guide local collapsing from nearby variants in the functional domain.

Specifically, POINT incorporates the 3D protein structure into the kernel machine regression framework, defining a local kernel function to enable variant-specific information collapsing. For a given variant, the amount of information contributed from its neighboring variants decays with the distance between variants in the 3D protein space. POINT performs local score tests for each variant over a range of kernel scale values, adaptively choosing the maximum distance allowed for information collapsing. In particular, for each variant, POINT calculates the minimum p-value (minP) across different distances, and uses a resampling approach to compute the p-value of minP, which can then be used to rank and select promising variants.

Below we evaluate the prioritization performance of POINT using simulation studies. We also apply POINT to the Action to Control Cardiovascular Risk in Diabetes (ACCORD) clinical trial data, finding promising rare variants in *PCSK9* that may explain variations in low-density lipoprotein (LDL) level.

## Methods

### Overview of POINT

We consider a study of *n* subjects with phenotype *Y*_*n×*1_ = [*Y*_1_, *…, Y*_*n*_]^*T*^. We assume *Y* follows an exponential family distribution with canonical link *g*(*µ*) = *g*(*E*[*Y |X, G*]), where *G*_*n×M*_ is the genotype design matrix of the *M* variants, and *X*_*n×p*_ is a matrix of the *p* non-genetic covariates. A kernel machine (KM) model for the local effect of variant *m, m* = 1, *…, M*, is of the form

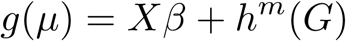

where 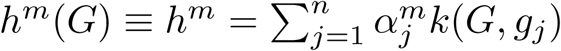 is a *n ×* 1 vector of the effect of variant *m*, and 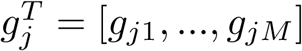 is the *j*th row of *G* and is genotype design vector for individual *j*. We assume 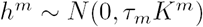, where the *n × n* matrix *K*^*m*^ = *{k*^*m*^(*g*_*i*_, *g*_*j*_)*}* is a local kernel matrix for variant *m*, describing the covariance between the local effect of variant *m* for different individuals. The local kernel matrix *K*^*m*^ is constructed in a manner such that *K*^*m*^ only puts non-trivial weights on the genetic similarity from variants that are in close proximity to variant *m*, with closer neighboring variants receiving higher weights. As detailed later, the local kernel function *k*^*m*^(*g*_*i*_, *g*_*j*_) uses the distance between variants in the 3D protein space to determine the amount of contribution from neighboring variants when quantifying the localized genetic similarity about variant *m*. From the local kernel, we construct a local kernel test with null hypothesis *H*_0_ : *τ*_*m*_ = 0 to evaluate if variant *m*, along with its proximal neighboring variants, are associated with the phenotype.

POINT consists of five main steps: (1) obtain the position of each variant in the 3D protein space, (2) construct a variant correlation matrix using the Euclidean distance between variants in the 3D protein space, (3) construct protein structure guided kernel matrices, (4) perform a local kernel test of *H*_0_ : *τ*_*m*_ = 0 for variant *m* over a range of collapsing distances and obtain the p-value, and finally (5) perform post hoc annotations of identified variants. The workflow is illustrated in Fig 1. Each step is further described below.

**Fig 1.**
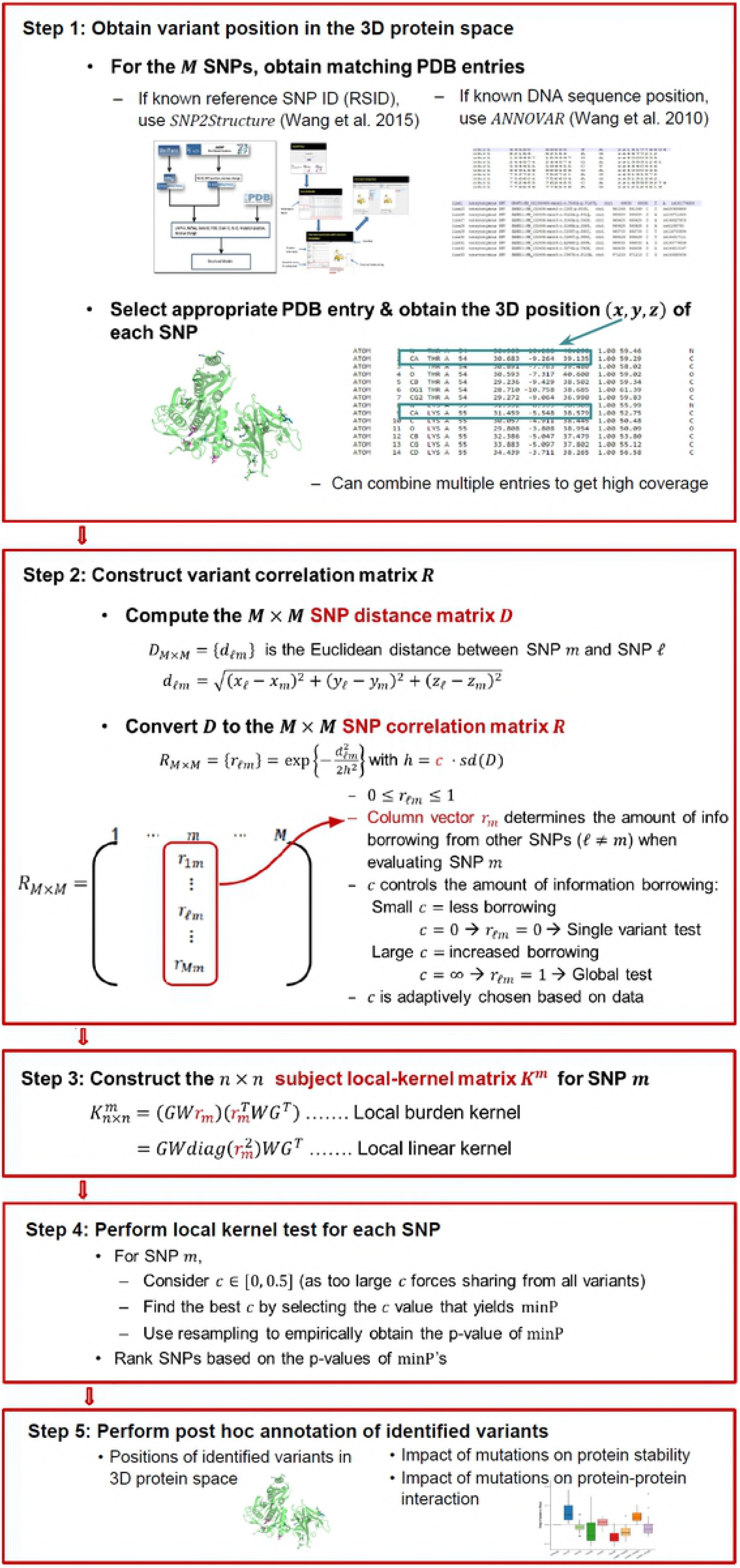
Overview of the protein structure guided local test (POINT).

#### Step 1: Obtaining Variant Positions in the 3D Protein Space

The first step in performing the protein structure-guided local test is to collect protein tertiary structure information, in the form of 3D coordinates in the protein space, for each of the *M* variants of interest. In order to do so, we must first map our genotype data to an appropriate Protein Data Bank (PDB) [32] entry. When the reference SNP ID (RSID) is known for each variant, the structural database online query tool SNP2Structure [33] is used to directly obtain a mapping of each SNP to their corresponding amino acid position for all relevant PDB entries. When only the variant position on the DNA sequence is known, we use the annotation tool ANNOVAR [34] to obtain the gene name, DNA mutation, and corresponding amino acid position and mutation of each SNP of interest. These amino acid mutations can be manually aligned to PDB entries for the gene of interest.

When a gene has multiple protein structure entries available on PDB, we select an appropriate entry for our analysis. Optimal PDB structures should have high resolution (*≤* 2.0Å), good data quality (e.g., low percent outliers, clashscore and Rfree score), and high coverage of variant 3D protein position for our variant set. Once an appropriate PDB entry has been identified, we extract the 3D coordinates (*x, y, z*) for each variant of interest, either using the coordinates of the carbon alpha for that particular amino acid residue, or taking the average of the coordinates of all the atoms forming the side chain of that residue (also called side chain centroid).

#### Step 2: Constructing Variant Correlation Matrix *R*

From the 3D Cartesian coordinates obtained in Step 1, we build a SNP pairwise distance matrix, *D*_*M×M*_ = *{d*_*ℓm*_*}*, where *d*_*ℓm*_ =[(*x*_*ℓ*_ *- x*_*m*_)^2^ + (*y*_*ℓ*_ *- y*_*m*_)^2^+(*z*_*ℓ*_ *- z*_*m*_)^2^]^1^*/*^2^ is the Euclidean distance between variants *ℓ* and *m* on the protein tertiary structure, and *d* _*ℓm*_ = 0 if *ℓ* = *m*. Using distance matrix *D*, we form a *M × M* variant correlation matrix 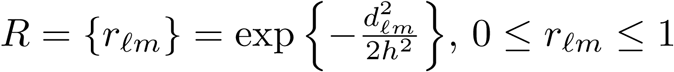.

Although we call *R* the variant correlation matrix, rather than inducing correlation between variants, matrix *R* is used to induce smoothing of information between neighbors, allowing gradual drop-off in the amount of borrowing between variants as the variant distance increases. Specifically, let *r*_*m*_ be the *m*^*th*^ column vector of *R* with dimension *M ×* 1; when testing for variant *m, r*_*m*_ determines the relative contribution from each other variant *ℓ* = 1, *…, M* (noting that *r*_*mm*_ = 1) via scale parameter *h*.

Rather than performing parameter tuning to determine an optimal scale *h*, we follow a similar approach to Tango [35] and Schaid et al. [27] and examine a grid of values. However, to make *h* scale free, instead of using proportion of maximum distance as a metric, we consider a grid over the proportion of the standard deviation of all pairwise Euclidean distances between variants, i.e., *h* = *c · sd*(*D*). The idea behind using a function of the standard deviation of pairwise distances is borrowed from nonparametric theory for choosing the optimal bandwidth of kernels.

Expressed in this manner, parameter *c* is used as a proxy for separating variants, so that only those variants within the “neighborhood” (or cluster) of variant *m* on the protein structure are likely to contribute information when quantifying localized genetic similarity around variant *m*. Larger *c* values encourage information contributed from a larger neighborhood, and the local test becomes a global-level test (i.e., *r*_*ℓm*_ = 1 for all *ℓ*) when *c* = *∞*. Smaller *c* values allow information to be contributed from a smaller neighborhood, and the local test becomes a single-variant test (i.e., *r*_*mm*_ = 1 and *r*_*ℓm*_ = 0 for *ℓ ≠m*) when *c* = 0.

When conducting local kernel tests in Step 4, we consider a grid of *c* values between 0 and 0.5 and let the data adaptively choose the best scale *c*. This strategy provides two layers of protection to safeguard against false positive selections. First, as illustrated later in Table 1 of simulation results, setting *c* = 0.5 as the maximum proportion of standard deviation of distance allowed for borrowing forces information sharing only from variants within a localized neighborhood in the protein tertiary space. In contrast, a larger maximum *c* value would result in information sharing from variants across the protein, regardless of whether they are close enough to be expected to share biological architecture or not. Consequently, a larger maximum *c* may lead to higher chances of selecting non-causal variants as promising loci. The second layer of protection stems from the fact that with the adaptively determined *c*, structure is used as a prior where, rather than forcing sharing of information between variants which may not have related genetic effects on phenotype, neighboring information can be shared only if there appears to be sufficient support from the data to do so.

**Table 1.**
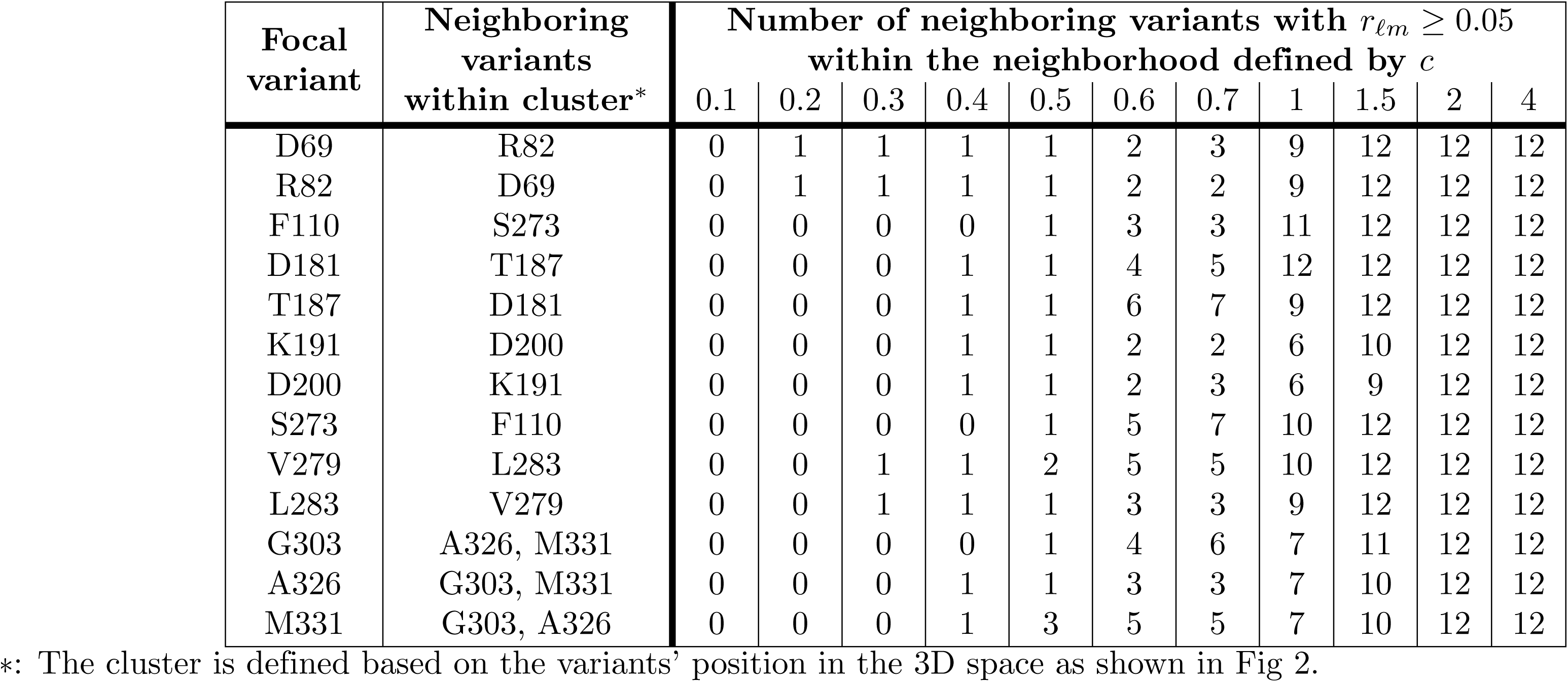
Counts of neighboring variants which contribute “significantly” to the focal rare variants in *PLA2G7*. Neighboring variants *ℓ*’s contributing *≥* 5% of the information to the focal variant *m* (i.e., neighboring variants with *r*_*ℓm*_ *≥* 0.05) for different values of *c*.

| Focal variant | Neighboring variants within cluster* | Number of neighboring variants with $r_{\ell m} \geq 0.05$ within the neighborhood defined by $c$ | | | | | | | | | | |
| --- | --- | --- | --- | --- | --- | --- | --- | --- | --- | --- | --- | --- |
|  |  | 0.1 | 0.2 | 0.3 | 0.4 | 0.5 | 0.6 | 0.7 | 1 | 1.5 | 2 | 4 |
| D69 | R82 | 0 | 1 | 1 | 1 | 1 | 2 | 3 | 9 | 12 | 12 | 12 |
| R82 | D69 | 0 | 1 | 1 | 1 | 1 | 2 | 2 | 9 | 12 | 12 | 12 |
| F110 | S273 | 0 | 0 | 0 | 0 | 1 | 3 | 3 | 11 | 12 | 12 | 12 |
| D181 | T187 | 0 | 0 | 0 | 1 | 1 | 4 | 5 | 12 | 12 | 12 | 12 |
| T187 | D181 | 0 | 0 | 0 | 1 | 1 | 6 | 7 | 9 | 12 | 12 | 12 |
| K191 | D200 | 0 | 0 | 0 | 1 | 1 | 2 | 2 | 6 | 10 | 12 | 12 |
| D200 | K191 | 0 | 0 | 0 | 1 | 1 | 2 | 3 | 6 | 9 | 12 | 12 |
| S273 | F110 | 0 | 0 | 0 | 0 | 1 | 5 | 7 | 10 | 12 | 12 | 12 |
| V279 | L283 | 0 | 0 | 1 | 1 | 2 | 5 | 5 | 10 | 12 | 12 | 12 |
| L283 | V279 | 0 | 0 | 1 | 1 | 1 | 3 | 3 | 9 | 12 | 12 | 12 |
| G303 | A326, M331 | 0 | 0 | 0 | 0 | 1 | 4 | 6 | 7 | 11 | 12 | 12 |
| A326 | G303, M331 | 0 | 0 | 0 | 1 | 1 | 3 | 3 | 7 | 10 | 12 | 12 |
| M331 | G303, A326 | 0 | 0 | 0 | 1 | 3 | 5 | 5 | 7 | 10 | 12 | 12 |

\*: The cluster is defined based on the variants’ position in the 3D space as shown in Fig 2.

#### Step 3: Constructing Local Subject Kernel Matrix *K*^*m*^ **for Variant** *m*

Given the *M × M* variant correlation matrix *R*, we create the *n × n* subject kernel matrix *K*^*m*^ that quantifies the genetic similarity between all pairs of individuals at variant *m* and its neighboring variants. By incorporating information from *R* as an additional weight, the commonly used global kernel functions can be extended to local kernels, where genetic similarity is calculated based largely on focal variant *m* and less on neighboring variants, with effect decaying with distance. To illustrate, let *wℓ* be the variant-specific weight for variant *ℓ* (e.g., based on the MAF or functional impact of variant *ℓ*) and let *r*_*ℓm*_ be the (*ℓ, m*)^*th*^ entry of the variant correlation matrix *R*. To quantify the similarity between subjects *i* and *j*, the global burden kernel function is 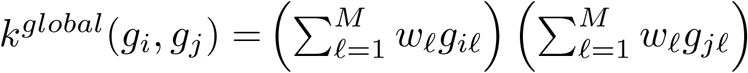 [36]. The local burden kernel function can be obtained as 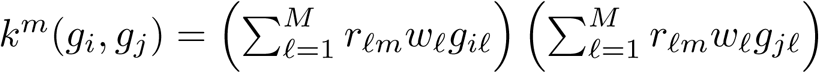. The additional weight *r*_*ℓm*_ in the local kernel function controls the amount of contribution from variant *ℓ*, which diminishes as variant *ℓ*’s distance from the focal variant *m* increases. Following this idea and using a matrix representation, we have the local burden kernel as 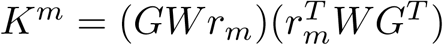 where *W* = *diag*(*w*_1_, *…, w*_*M*_), the linear local kernel as 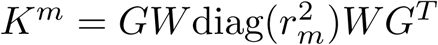, and the polynomial local kernel as 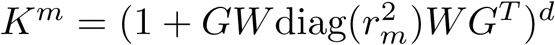.When *r*_*m*_ = **1**_*M×*1_, the local kernel matrix becomes the global kernel matrix.

#### Step 4: Performing Local Kernel Test of *H*_0_ : *τ*_*m*_ = 0 **for Variant** *m*

The local test of *H*_0_ : *τ*_*m*_ = 0 assesses whether variant *m*, along with its nearby variants (within a small neighborhood defined by *c*) are associated with the phenotype. To describe the score-based test statistic of the local test, we further rewrite the kernel matrix *K*^*m*^ as *K*^*m,c*^ to emphasize that it is computed at a fixed value of *c*. Following Wu et al. [8] and Tzeng et al. [37, 38], the score-based test statistic *T*_*m,c*_ has a quadratic form and follows a weighted chi-square distribution asymptotically. Specifically, 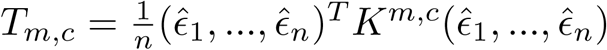, where 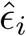 is the fitted residual for the KM model under the null hypothesis, i.e., 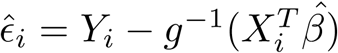 where 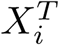 is the 1 *× p* row vector of the covariate design matrix *X*. The weights of the weighted chi-square distribution are given in the Appendix. Therefore, for a fixed *c*, the corresponding p-value, denoted by *p*_*m,c*_, can be calculated using the Davies method [39].

Given a grid of *c*’s, *c* = *c*_1_, *…, c*_*L*_, we adaptively find the optimal *c* by choosing the value that yields the minimum *p*-value for variant *m*, i.e., min 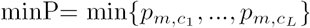. As shown in the Appendix, we develop a resampling approach to calculate the p-value of the minP statistic, denoted by 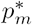. These p-values can be used to rank and select promising variants, e.g., to select the top variants whose p-value 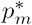 is less than a certain threshold.

#### Step 5: Performing Post Hoc Annotations of Identified Variants

Once the promising variants are identified, we examine the functional and structural consequences of these mutations. The post-hoc examinations include the qualitative examination of the variant locations on the protein three-dimensional structure. Moreover, complementary cheminformatics calculations, such as molecular dynamic simulations (MDS), are used to predict how the identified mutations could affect the protein flexibility and overall stability (which in turn could affect the protein-protein interactions and the overall phenotype of interest). The full-atom simulations enable us to estimate some protein’s conformational changes of atomic positions via root mean square deviation (RMSD) and per-residue root mean square fluctuation (RMSF), compared to the wildtype protein structure.

### Simulation Study Set-Up

We design a simulation following the work of Song et al. [23], which examined the effect of SNPs within Phospholipase A2 Group VII (*PLA2G7*) on protein function and enzyme activity of Lipoprotein-associated phospholipase A_2_ (Lp-PLA2) measured on *∼*90 individuals. Genotype data from Sanger sequencing of *PLA2G7* are also available on 2000 individuals from the CoLaus study, a study examining psychiatric, cardiovascular, and metabolic disorders in 6188 Caucasians aged 35-75 from Lausanne, Switzerland [40, 41]. Song et al. [23] found that variants which are deemed likely to be non-null variants for enzyme activity of Lp-PLA2 tend to cluster together and are predominately on the surface of protein, while null variants are nearby in the core of protein [23]. For the simulation study, we obtain the sequencing genotypes of *PLA2G7* from Song et al. [23] and obtain the variants’ 3D coordinates on the protein tertiary structure from PDB entry 3F96 [42]. In total, 13 rare variants from the Song et al. [23] study have protein coordinate information available in PDB; their variant information is provided in S1 Table. Fig 2 shows the variants’ location in the 3D protein structure and the corresponding Euclidean distance-based clustering of these variants. In the figure, each variant is named by its amino acid position on the 2D sequence structure.

**Fig 2.**
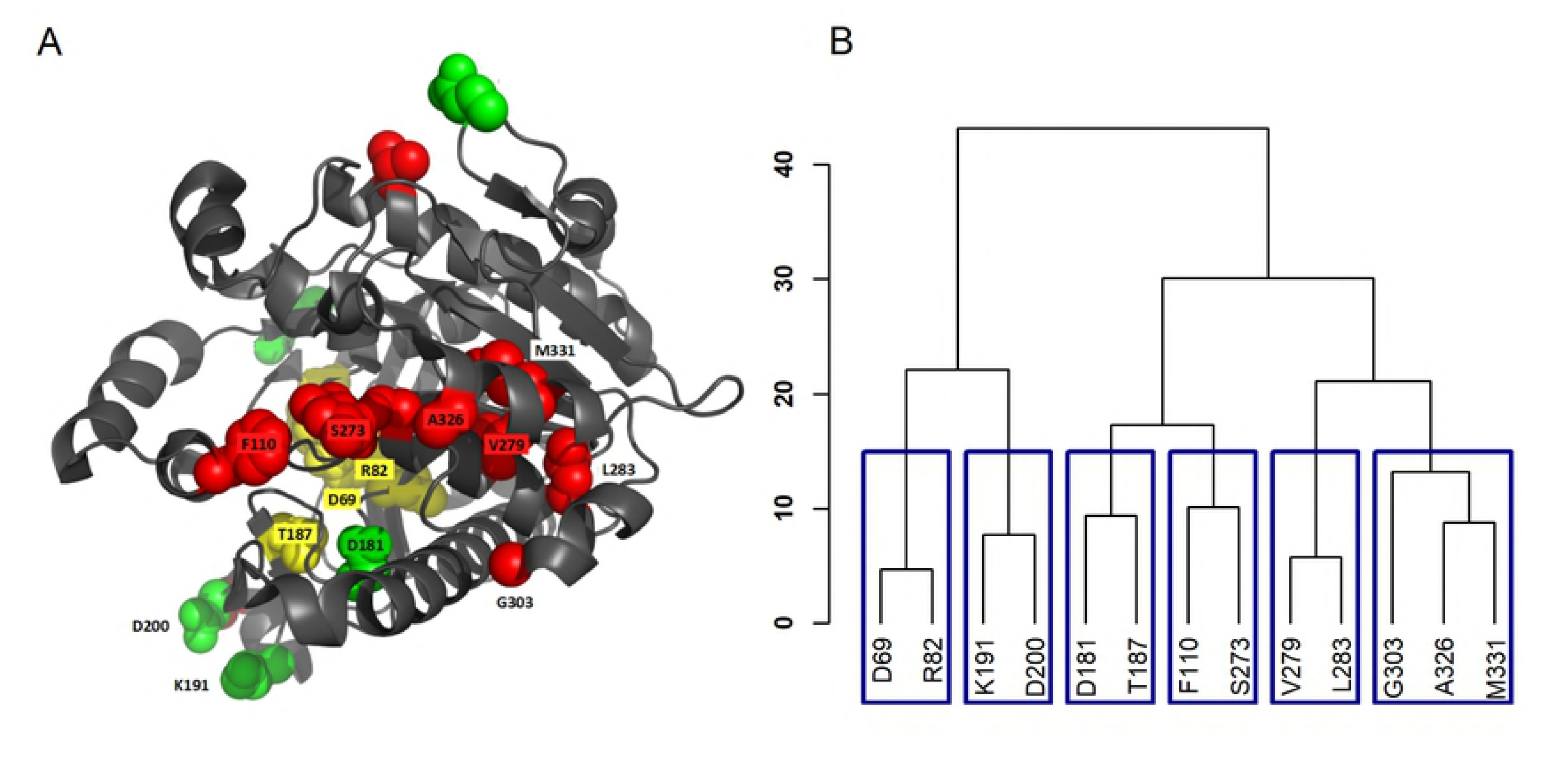
*PLA2G7* rare variant positions. A: Rare variant locations on the protein tertiary structure. B: Corresponding Euclidean distance-based clustering of the variants.

Using the genotype data from these 13 variants, we generate phenotypes for 1,000 individuals from the model of *g*(*µ*) = *β*_0_ + *Gβ*_*G*_, with identity link *g*(*µ*) = *µ* for continuous traits (i.e., 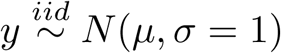), and logit link *g*(*µ*) = exp(*µ*)*/*(1 + exp(*µ*)) for binary traits. We set the intercept *β*_0_ = 0.5 for continuous traits, and *β*_0_ = *-*0.05 for binary traits, and set the coefficient vector ***β****G* = *{β*_*G,m*_*}* of genetic effects as *β*_*G,m*_ = *b × |* log_10_(*MAF*_*m*_)*|*, where *b ≠*0 for causal variants and is equal to zero otherwise, and *MAF*_*m*_ is the minor allele frequency of variant *m*. This specification of *β*_*G,m*_ assigns larger effects to rarer variants.

We consider a variety of scenarios for causal variants: Scenario (A): One cluster is causal, where the causal variants cluster close together on the tertiary protein structure, with varying closeness on the amino acid sequence; we consider four sub-scenarios with (D69, R82), (G303, A326, M331), (K191, D200), and (P110, S273) chosen to be the causal variant clusters. Scenario (B): Part of a cluster is causal, where only a subset (e.g., two of three) of closely clustered variants are causal; we consider two sub-scenarios where (A326, M331) and (G303, A326) are causal from the variant cluster (G303, A326, M331). Scenario (C): Two opposing clusters are causal, where two clusters of variants are causal, with one cluster, (D69, R82), positively conferring phenotype risk, and another cluster, (G303, A326, M331), negatively conferring phenotype risk. We also consider the scenario of no causal variants to examine the validity of the proposed POINT tests.

For POINT, we use weights proportional to a Beta(MAF,1,25) distribution as described in Wu et al. [8] (i.e., *w*_*ℓ*_= (1 *- MAF*_*ℓ*_)^24^) to upweight the contribution of rare neighboring variants. We consider a grid of 6 values for *c*, i.e., *c* = (0, 0.1, 0.2, 0.3, 0.4, 0.5) and perform tests using burden and linear kernels, each with 500 replications per scenario, and 1000 resamples per replication. We evaluate the ability of POINT to prioritize causal variants by comparing to the single variant score test (which corresponds to POINT with *c* = 0 and is referred to as SVT) and the scan statistic of Ionita-Laza et al. [25] (which is referred to as SCAN). SCAN has been shown to be the superior method among those methods searching for genomic clustering of risk variants [27]. However, SCAN is only used as a comparison in the binary case-control simulations, as it is not directly applicable to continuous phenotype data.

The selection performance of the three methods is assessed using true positive rates (TPR), false discovery rates (FDR), and a composite metric based on the F measure, which is the harmonic mean of the TPR and FDR. TPR is obtained by first computing the fraction of selected causal variants among all causal variants in each replication, and then averaging across the 500 replications. FDR is obtained by first computing the fraction of selected non-causal variants among all selected variants in each replication, and then averaging across the 500 replications. For SVT and POINT, a variant is selected if its p-value is smaller than a pre-specified threshold, e.g., 0.05. For SCAN, a variant is selected if it is included in the best window (i.e., the window with maximum test statistic) and the best window is significant. We also evaluate the overall selection performance using empirical receiver operating characteristic (ROC) curves to show the results across all possible decision (e.g., p-value) thresholds.

### Application to the ACCORD Study and ***PCSK9***

The ACCORD clinical trial was a multi-center trial with the intent to test for the effectiveness of intensive glycemic, blood pressure, and fenofibrate treatments versus their corresponding standard treatment strategies on cardiovascular disease (CVD) endpoints in subjects with type 2 diabetes [43–46]. The trial enrolled 10,251 subjects with type 2 diabetes and a risk or history of CVD from 77 centers around North America, and found that intensive treatments were not beneficial and even potentially harmful for some of the CVD endpoints studied [44]. A recent study of this trial investigated genotype associations with individual variation in serum lipid levels in the context of patients with type 2 diabetes [46]. Focusing on the baseline pre-intervention data, Marvel and Rotroff et al. [46] examined the association between baseline blood lipid levels and common variants and rare variants from 16,538 genes in 7844 ACCORD trial participants that consented to genetic studies. Based on rare variant associations, they found 11 genes to be significantly associated with blood lipid levels, including total cholesterol, LDL, high-density lipoprotein, and total triglycerides.

Here we focus on proprotein convertase subtilisin/kexin 9 (*PCSK9*), as it is the gene reported to be most highly associated with LDL from the baseline study of Marvel and Rotroff et al. [46] and of high clinical importance. Because the gene-level rare variant signals in Marvel and Rotroff et al. [46] were mainly identified via burden-based tests, we apply POINT with burden kernels, aiming to prioritize the individual variants associated with LDL within *PCSK9*. Following the work of Marvel and Rotroff et al. [46], we considered rare variants to be those with MAF *<* 3% and use only individuals with less than 15% missingness. Missing genotype information was imputed previously by Marvel and Rotroff et al. [46]. We use the IBCI database SNP2Structure [33] to match each variant RSID with their corresponding amino acid position on the 2D sequence, as well as the protein chains that the variants are situated on in all PDB entries for *PCSK9*. We obtain carbon alpha coordinates from PDB entry 4K8R [47], which we determined to be the most representative of the wild type protein while maximizing the number of variants of interest with known protein tertiary position, i.e., 19 of 22 variants. A summary of the 19 variants and the corresponding 3D coordinates from PDB is given in S2 Table. In the analysis, we adjust for 26 baseline covariates as in Marvel and Rotroff et al. [46], including patient age, gender, body mass index (BMI), presence of cardiovascular history, trial treatment arm assignment, top three principal components of ethnic background, years since diabetes and since hyperlipidemia diagnoses, fasting glucose level, and indicators of use of different treatments (e.g., insulin, lipid-lowering drugs, etc.). A full list of these covariates can be found in the Supplementary Materials of Marvel and Rotroff et al. [46].

To better understand the biological significance of the promising variants identified, we further examine whether the mutant sequences could have significant impact on the PCSK9-LDLR binding stability compared to the wildtype sequence. Using MDS with 3 replicates for each sequence, we examine the conformational changes of mutant proteins. We first measure the atomic mobility via RMSF for the wildtype protein and for each of the PCSK9 mutant proteins. We then quantify the stability changes for the PCSK9-LDLR interaction by calculating the RMSF difference between the mutant and the wildtype, and use the Wilcoxon rank-sum test to detect any significant differences.

## Results

### Results of Simulation Studies

#### Variant correlation matrix *R* vs. information borrowing from other variants

To illustrate how the variant correlation matrix *R* controls information borrowed from nearby variants, we examine how the range of variants that a focal variant significantly borrows from changes for different values of *c*. For each variant in *PLA2G7*, Table 1 lists its neighboring variant(s) within the same cluster given their position on the 3D protein space (as seen in Fig 2), and the number of variants (excluding the focal variant itself) that “significantly” contribute information to the focal variant for a range of *c* values between 0.1 and 4. Here we use the term “significantly” to loosely indicate those variants *ℓ* whose value *r* _*m*_ is at least 0.05, i.e., contributing at least 5% as much information as the focal variant. We see that for *c* = 0.5, the number of variants that contribute information to the focal variant corresponds well with the true number of variants within the same cluster as it. As *c* increases past 0.5, the number of variants borrowed from for each variant increases dramatically—borrowing from many more than just those that cluster together even for *c* = 0.6 and *c* = 0.7, and borrowing from all variants as *c* increases further.

In Fig 3 we show the relative 3D positions of the 13 rare variants in *PLA2G7*, focusing on variant V279 (highlighted in green). Each other variant is colored to indicate the magnitude of contribution into the local genetic similarity for V279 over different values of *c*, with those colored in lighter pink having lower contribution, and darker pink indicating higher contribution. In agreement with Table 1, we see that when *c* = 0.1, V279 does not significantly borrow information from any neighbors. When *c* = 0.3, it begins borrowing information from its closest neighboring variant, L283 (*r*_*L*283,*V* 279_ *≈* 0.2). For *c* = 0.5, this amount increases (*r*_*L*283,*V* 279_ *≈* 0.5), and we also see marginal contributions from M331 (*r*_*M*331,*V* 279_ *≈* 0.1). As *c* increases beyond 0.5 (e.g., *c* = 0.7), V279 begins to borrow information from variants spaced further away on the protein. For larger values of *c* (e.g., *c ≥* 2), the local test approaches a global test using similar weights for all variants along the protein. Similar patterns are observed when other variants are set as focal variants, as shown in S1 Fig - S12 Fig.

**Fig 3.**
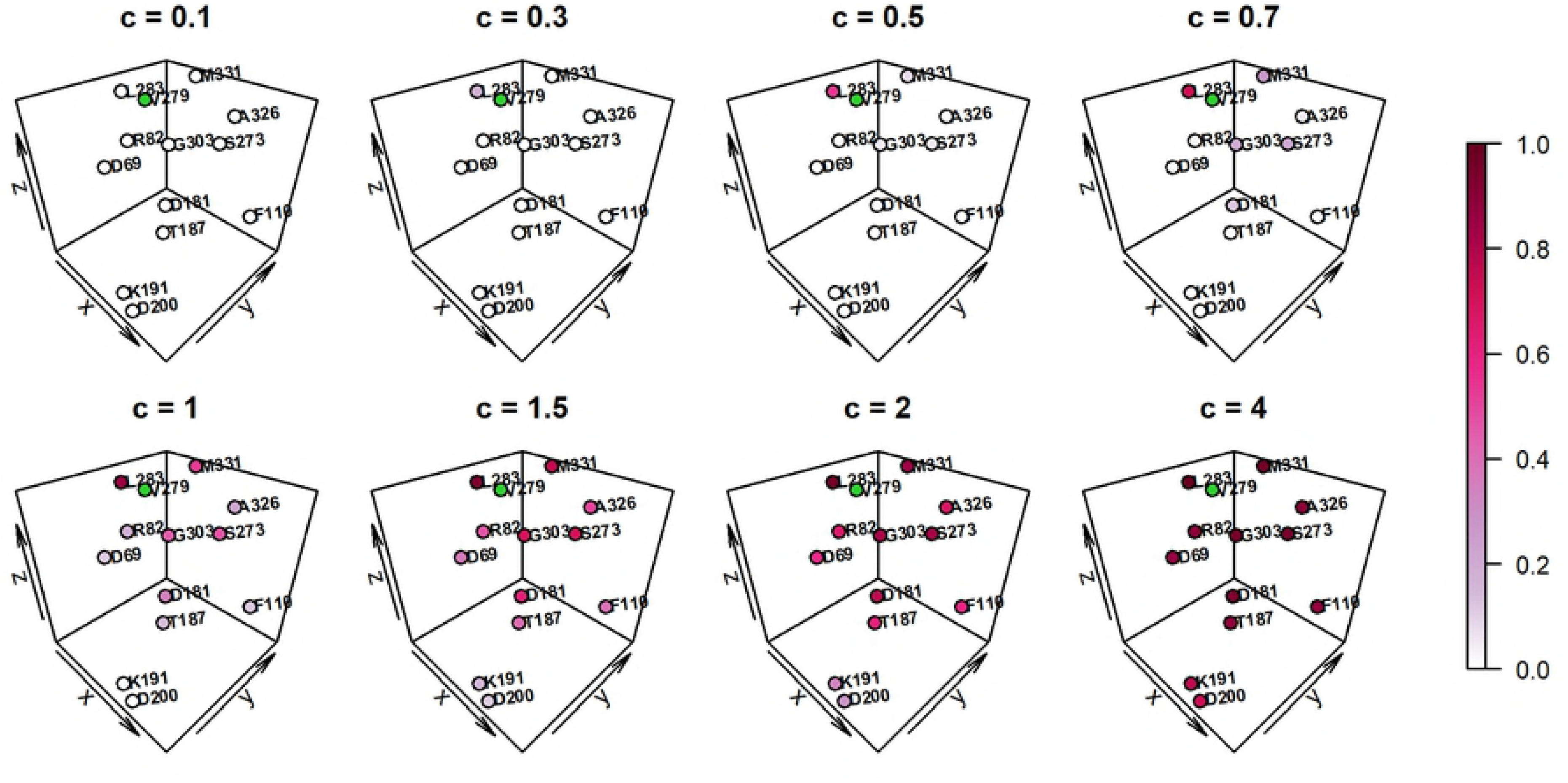
*PLA2G7* variant V279 borrowing map. Variant-borrowing map defining the amount of borrowing from neighboring variants for *PLA2G7* variant V279 for different values of *c*, with darker color representing higher levels of contribution via the variant correlation matrix *R*.

#### Selection performance of risk variants

To evaluate the performance of POINT, we first examine how it behaves under the null hypothesis of no causal variants. S13 Fig shows the quantile-quantile plots of the p-values for SVT, POINT (applied using burden and linear local kernels), and SCAN methods for both quantitative traits (top) and binary traits (bottom). The quantile-quantile plots compare the observed p-values with the expected p-values under the null. For all methods, the points fall near the 45 degree line, confirming the validity and appropriate implementation of each method.

We summarize the selection performance of SVT, POINT (again, with burden and linear local kernels, denoted by POINT-Burden and POINT-Linear, respectively), and SCAN under each causal scenario for a p-value threshold of 0.05 in Tables 2 (for continuous traits) and 3 (for binary traits). In Table 2, the relative true positive and false discovery rates of the methods show that for continuous traits, (a) POINT-Burden and POINT-Linear have similar or better selection performance than SVT, and (b) POINT-Burden has better or similar performance compared to POINT-Linear. These results are as expected: for (a), POINT includes SVT as a special case; therefore when the data do not support borrowing information from neighbors (e.g., in Scenario B2, where causal variant A326 has its nearest neighbor M331 as non-causal), POINT tends to adaptively set *c* = 0 and perform a single variant test. For (b), the effect mechanisms used to simulate trait values in the simulation are such that local burden collapsing would be the most efficient kernel to capture the association effects.

**Table 2.** Continuous trait simulation selection performance. Selection performance for single variant test (SVT), POINT test using local burden kernel (POINT-Burden), and POINT test using local linear kernel (POINT-Linear) for continuous trait simulation study

| Causal Scenario | Causal Variants | True Positive Rate |  |  | False Discovery Rate |  |  | F measure |  |  |
| --- | --- | --- | --- | --- | --- | --- | --- | --- | --- | --- |
|  |  | SVT | Burden | POINT<br>Linear | SVT | Burden | POINT<br>Linear | SVT | Burden | POINT<br>Linear |
| (A)<br>One Causal<br>Cluster | A1 (D69, R82) | 0.736 | 0.971 | 0.951 | 0.435 | 0.391 | 0.373 | 0.639 | 0.749 | 0.756 |
|  | A2 (G303, A326, M331) | 0.802 | 0.930 | 0.839 | 0.177 | 0.159 | 0.163 | 0.813 | 0.883 | 0.838 |
|  | A3 (K191, D200) | 0.589 | 0.800 | 0.648 | 0.302 | 0.237 | 0.276 | 0.639 | 0.781 | 0.684 |
|  | A4 (F110, S273) | 0.730 | 0.844 | 0.797 | 0.261 | 0.234 | 0.233 | 0.734 | 0.803 | 0.782 |
| (B)<br>Part of Cluster<br>is Causal | B1 (A326, M331) | 0.778 | 0.906 | 0.810 | 0.246 | 0.217 | 0.232 | 0.766 | 0.840 | 0.788 |
|  | B2 (G303, M326) | 0.870 | 0.882 | 0.863 | 0.242 | 0.249 | 0.239 | 0.810 | 0.811 | 0.809 |
| (C)<br>Two Opposing<br>Causal Clusters | C<br>(D69, R82) deleterious &<br>(G303, A326, M331) protective | 0.823 | 0.971 | 0.925 | 0.207 | 0.205 | 0.195 | 0.808 | 0.874 | 0.861 |

We observe a similar pattern of relative performance among SVT, POINT-Burden and POINT-Linear in the binary trait simulations (Table 3) as in continuous traits. For binary traits, we also examine the relative performance of SCAN, and see that, while SCAN often has good performance, its overall performance depends largely on whether or not the causal variants cluster together on the 2D sequence. When the causal variants are not sequential on the 2D sequence (e.g., in Scenario A4 where (P110, S273) are causal, and in Scenario C where (D69, R82) and (G303, A326, M331) are causal), SCAN struggles to identify the true best window and gives sub-optimal selection results.

**Table 3.**
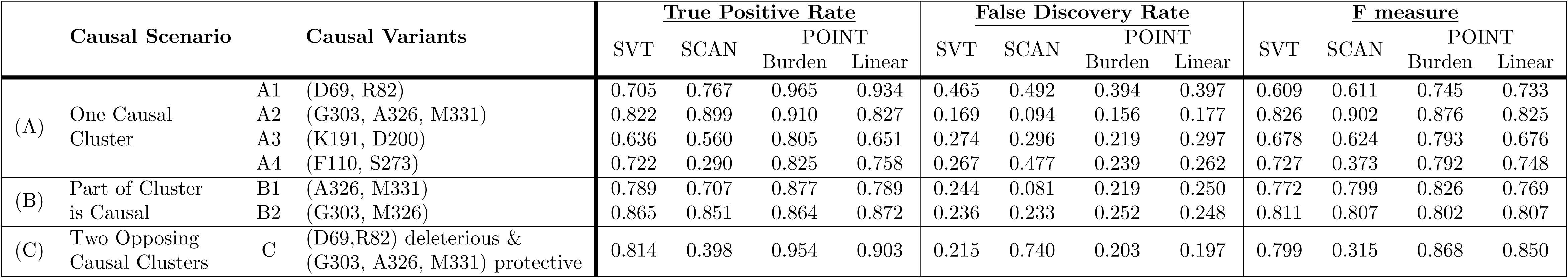
Binary trait simulation selection performance. Selection performance for single variant test (SVT), POINT test using local burden kernel (POINT-Burden), POINT test using local linear kernel (POINT-Linear), and scan test (SCAN) for binary trait simulation study

To complete the picture and to assure comparability among different methods, we also examine the relative selection performance of the methods over different selection criteria using ROC curves. The results are shown in S14 Fig (for continuous traits) and S15 Fig (for binary traits). The ROC plots evaluate the true positive rate of variant selection (sensitivity) of the test against the false positive rate of variant selection (1-specificity) over all possible ranges of selection thresholds. Better selection methods have larger area under the ROC curve, with plots approaching upper the upper left corner, where more causal variants are selected with fewer null variants selected. We observe a similar pattern of relative performance over the possible range of selection thresholds as is see in Tables 2 and 3.

In summary, POINT gives stable selection performance across various scenarios, and performs at least as well as – often improving upon – the single variant analysis by yielding higher true positive rates across different decision thresholds.

### Results of Application to the ACCORD Study and ***PCSK9***

Table 4 shows the analysis results for *PCSK9* using SVT and POINT. We select those variants with p-value less than 0.05 as promising variants. Using this criterion, there are two promising variants identified using both of SVT and POINT: A443 and H553. POINT further identifies three additional variants: N157, H391, and N425. Fig 4A shows the variant positions on the 3D protein structure, with the two variants found by SVT and POINT in pink and red, and those found only by POINT in green and blue. Fig 4B shows the distance-based clustering of *PCSK9* variants based on their positions in the 3D protein structure.

**Table 4.** *PCSK9* analysis results summary. Results of *PCSK9* analysis using the single variant test (SVT) and POINT (POINT-Burden). Promising variants are selected using the criterion of p-value*<* 0.05 and are shown in bold font.

| Variant ID<br>(AA Coord.) | SNP RSID | SVT |  | POINT-Burden |  |  | Variants borrowed<br>information from* |
| --- | --- | --- | --- | --- | --- | --- | --- |
| | | p-value | Rank | Best $c$ | p-value | Rank | |
| R93 | rs151193009 | 0.159 | 7 | 0 | 0.195 | 11 | N/A |
| R96 | rs185392267 | 0.397 | 16 | 0.5 | 0.391 | 18 | R93 |
| **N157** | rs143117125 | 0.213 | 10 | 0.4 | **0.023** | 2 | H391, A443 |
| V252 | rs149139428 | 0.554 | 17 | 0.5 | 0.147 | 8 | L253, L283 |
| L253 | rs72646508 | 0.130 | 6 | 0.5 | 0.135 | 7 | V252 |
| P279 | rs72646509 | 0.318 | 13.5 | 0.4 | 0.408 | 19 | L283 |
| L283 | rs72646510 | 0.637 | 18 | 0.5 | 0.262 | 14 | V252, P279, H391 |
| **H391** | rs146471967 | 0.091 | 4 | 0.3 | **0.010** | 1 | N157, A443 |
| H417 | rs143275858 | 0.318 | 13.5 | 0.5 | 0.106 | 6 | N157, H391, A443, R525 |
| **N425** | rs28362261 | 0.051 | 3 | 0.5 | **0.034** | 4 | N157, A443 |
| **A443** | rs28362263 | **0.043** | 2 | 0.5 | **0.048** | 5 | N157, H391, H417, N425 |
| G466 | rs72646517 | 0.988 | 19 | 0.5 | 0.379 | 17 | R469, E498 |
| R469 | rs141502002 | 0.225 | 11 | 0.5 | 0.237 | 13 | G466, E498 |
| E498 | rs145468572 | 0.317 | 12 | 0.5 | 0.153 | 9 | G466, R469, R525 |
| R525 | rs140286279 | 0.347 | 15 | 0.4 | 0.374 | 16 | H417, E498 |
| **H553** | rs28362270 | **0.016** | 1 | 0 | **0.024** | 3 | N/A |
| Q554 | rs149311926 | 0.124 | 5 | 0 | 0.194 | 10 | N/A |
| Q619 | rs28362277 | 0.183 | 8 | 0 | 0.218 | 12 | N/A |
| R659 | rs147182054 | 0.201 | 9 | 0 | 0.359 | 15 | N/A |

\*: Only those variants with variant correlation matrix entry $r_{\ell m} \geq 0.05$ are listed.

**Fig 4.**
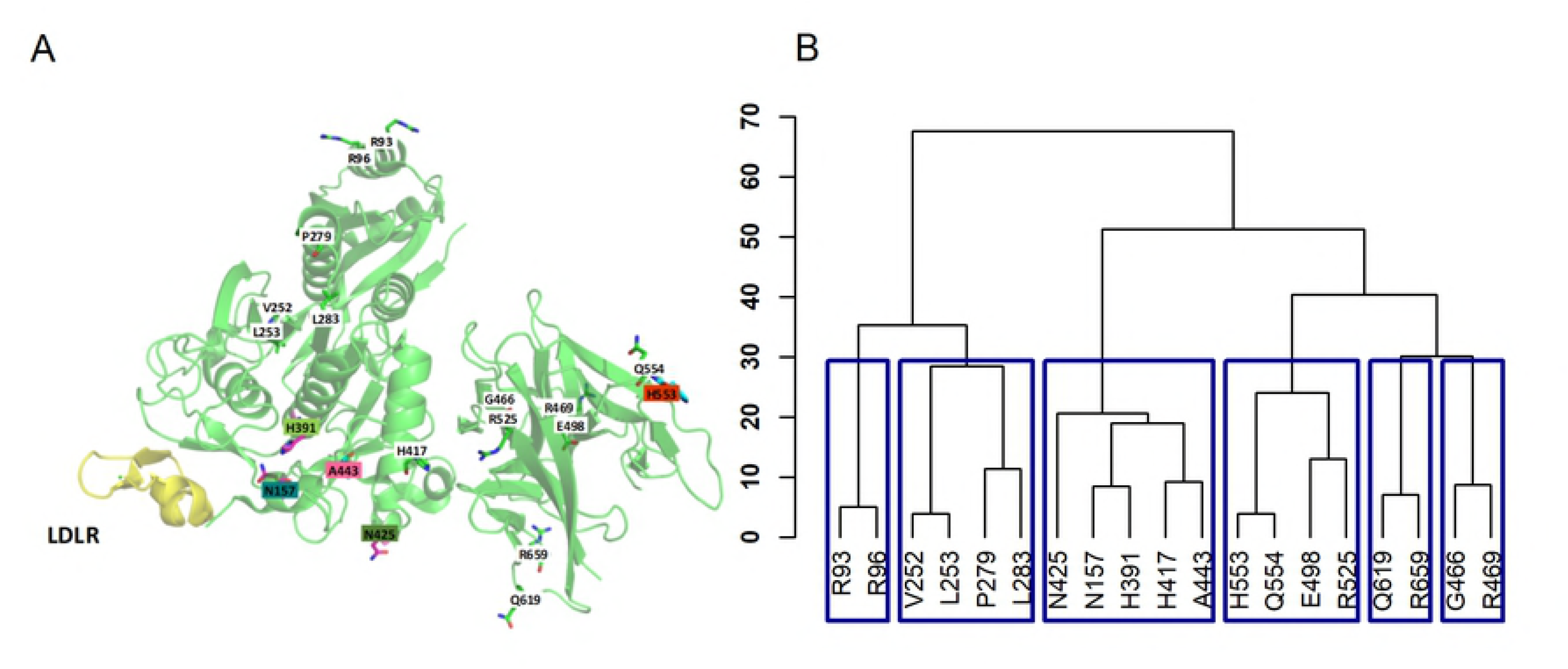
*PCSK9* rare variant positions. A: Rare variant locations on the protein tertiary structure of *PCSK9* binding with *LDLR* (shown in yellow). Promising variants (i.e., p-value*<*0.05) are shown in colored boxes, with variants found by both single variant test and POINT shown in red and variants found only by POINT shown in green and blue. B: Euclidean distance-based clustering of the variants.

S16 Fig shows the amount of information borrowed from neighboring variants for each variant at their chosen best value of *c*. We see that many of the promising variants identified by POINT cluster close together and choose to borrow information from one another. In particular, we see mutual borrowing of information between N157, H391, and A443, with A443 also borrowing from H417 and R525. The plots also show how information sharing between neighboring variants does not have to be symmetric. An example is H417, who chooses to borrow information from variants H391 and A443, though does not contribute to H391. The patterns of borrowing show how information sharing is variant-specific, allowing each variant to choose whether or not to borrow information based on how consistent the prior set by the local kernel is with the raw data. We further see that large association signal can occur without needing or choosing to borrow information from neighboring variants, as is the case with the selected variant H553, which chooses not to borrow from nearby variant Q554.

It has been shown that *PCKS9* impacts LDL levels by binding with *LDLR* (low-density lipoprotein receptor), prohibiting *LDLR* from binding LDL, and leading to increased LDL plasma levels [48]. As shown in Fig 4A, the variants newly identified by POINT were the closest variants in *PCSK9* to the protein-binding domain of *PCSK9* and *LDLR*. We also examine the MDS of the mutations from the promising loci. Based on the five variants identified using POINT, there are 8 unique mutant sequences observed in the ACCORD samples as listed in Fig 5. Importantly, 4 out of the 8 mutant sequences have significant conformational changes in protein-protein interaction when compared to the wildtype: single mutations N157K (p-value 2.38 *×* 10^−7^) and H553R (p-value 4.77 *×* 10^−7^), and double mutations A443T combined with H391N (p-value *×* 10^−5^) and A443T combined with N425S (p-value 1.67 *×* 10^−6^). The RMSF changes for the eight mutant sequences and corresponding p-values are shown in Fig 5. We note that a negative RMSF difference indicates that the amino acids involved in the protein-protein interaction have coordinates that fluctuate less than that of the wildtype (and hence the interaction is classically expected to be stronger). In contrast, a positive RMSF difference indicates that the amino acids move more than that of the wildtype (and hence the overall protein-protein interaction is expected to be weaker). Three of the significant sequences (i.e., N157, A443+H391, and A443+N425) are only identified by POINT. These results suggest a potential biological impact of these POINT-identified variants on the *PCSK9-LDLR* binding stability and hence an effect on the LDL level.

**Fig 5.**
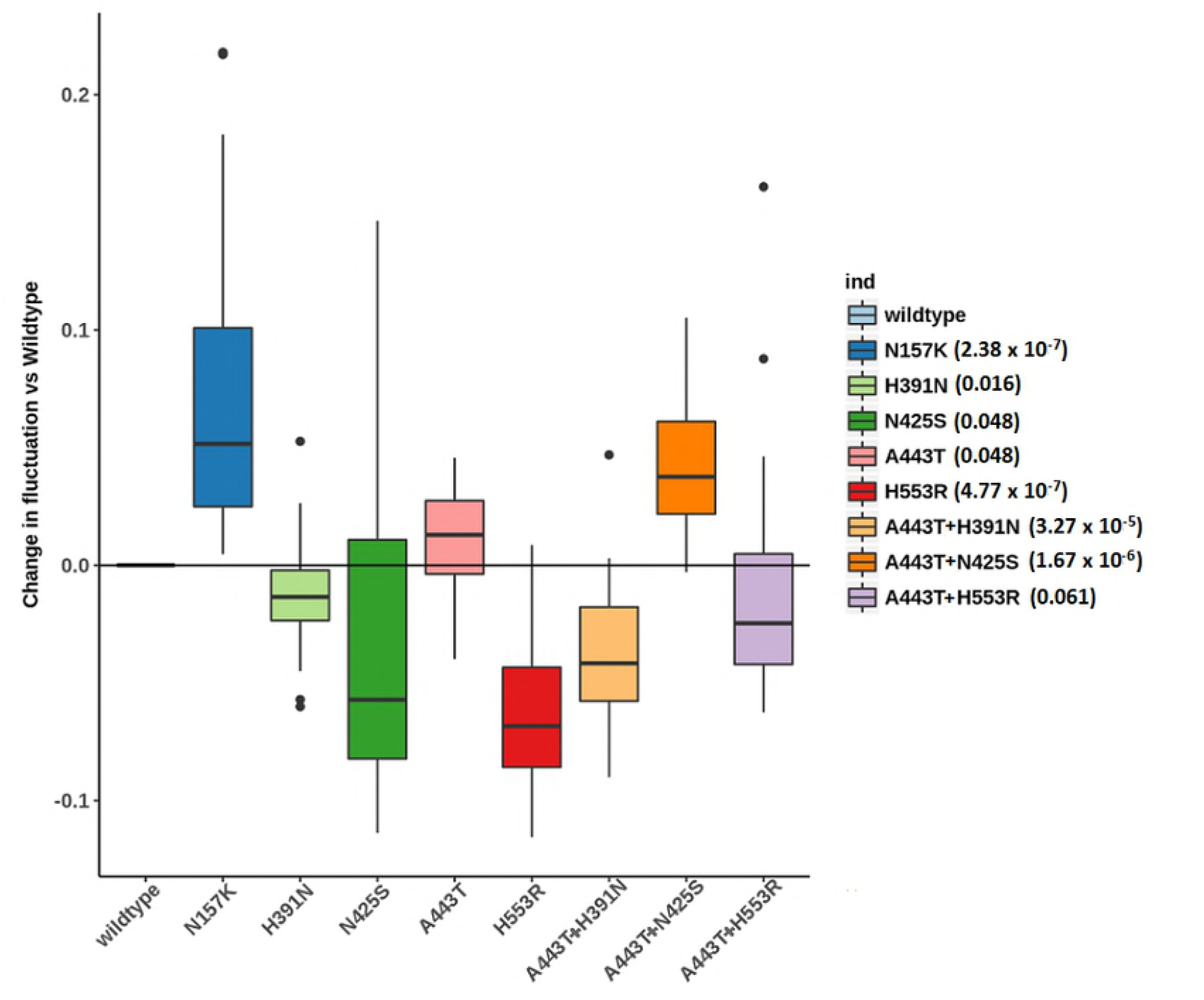
Assessment of *PCSK9-LDLR* binding stability for the mutant sequences from the five POINT-identified loci using molecular dynamic simulations. There are 8 mutant sequences observed in ACCORD samples, four of which have significant conformational fluctuation changes comparing to wildtype sequence: N157K, H553R, A443T+H391N, and A443T+N425S. P-values from Wilcoxon rank sum test of difference in RMSF in parentheses.

## Discussion

Here, we provide an analytic framework, POINT, to prioritize rare variants by incorporating protein 3D structure to guide local collapsing analysis. With POINT, we introduce a mathematical formulation of tertiary protein structure using a structural kernel, develop a statistical framework to perform inference at a localized level guided by the protein structure, and describe how the structure-supervised analysis can be used to identify variants likely to have an effect on the trait of interest. We have implemented the proposed analyses in R package POINT, available at http://www4.stat.ncsu.edu/˜sthollow/JYT/POINT/.

The performance of POINT is robust and stable across different scenarios investigated in this study. When the false positive rate is fixed at the same level, POINT has similar or improved ability to identify risk rare variants compared to alternate methods SVT and SCAN. POINT is adaptive, utilizing a data-driven scale *c* and the minimum *p* statistic to determine: (1) the appropriate neighboring variants to borrow information from, and (2) the optimal amount of information to borrow from those neighboring variants. As shown in the variant-borrowing maps (Fig 3, S1 Fig - S12 Fig, and Fig), while neighboring variants do tend to borrow from one another to gain strength, this borrowing only occurs when the data are supportive of the prior suggested by the protein structure and the borrowing does not have to be symmetric between a pair of variants.

Applying our local kernel method to the ACCORD clinical trial, we are able to pinpoint three new rare variants that are not found by single variant testing, all near the protein-binding domain between PCSK9 and LDLR. The results highlight the strength of our integrative method to find additional signal that cannot be found using single variant testing or MDS alone. This finding may have important clinical impact, given that PCSK9 inhibitors are a new class of drugs and are being accepted as a promising treatment for reducing LDL levels [49, 50].

POINT is constructed under the kernel machine framework with three main considerations that may affect performance: (1) choice of kernel, (2) choice of grid of *c* values, and (3) choice of PDB entry. For the first consideration, as noted in the literature, the local kernel test is valid even if a “wrong” kernel is chosen [9]. However, the power can be significantly affected by the choice of kernels because different kernel functions represent different underlying effect mechanisms (e.g., whether neighboring causal variants have similar or different effect patterns). Because such effect mechanisms are unknown a priori, choosing the “correct” or “optimal” kernel is still an important open problem in general kernel machine regression. One way to ensure the use of a “near optimal” kernel is to apply the composite kernel of Wu et al. [9], which can yield performance similar to the optimal kernel with substantial improvement over “wrong” kernels.

In our analyses we handle the second consideration by adaptively choosing a scale *c* over a grid ranging from *c* = 0 to *c* = 0.5. We show, using tables and variant borrowing maps, how the maximum *c* value affects how far from your focal variant you are willing to borrow information from. We choose *c* = 0.5 as our maximum grid value to ensure borrowing only from neighbors who may be considered to cluster close together on the protein tertiary surface, as the literature suggests common effects from closely clustered variants. As this is a multiplier of the standard deviation of distance between variants, this choice should also be applicable to different protein structures. Choosing a larger maximum *c* may be considered, but with caution so as not to increase false signal which may arise from borrowing outside of the cluster.

For the third consideration, we detail a few criteria for choosing an optimal protein structure entry from PDB, including good data quality and high coverage. In this work, we illustrate the POINT analysis under the scenario that the variants’ positions in the 3D protein structure can be obtained from a single PDB entry. However, in practice it is possible that no single entry has high coverage for the desired variant set. In this case, one can obtain the coordinate information by aligning multiple PDB entries with overlapping mapped residues using the PyMOL software (The PyMOL Molecular Graphics System, Version 1.8, Schrodinger, LLC.). When a variant in the set has no known coordinate information, instead of excluding it from the analysis as we did here, one may choose to include the variant by setting its Euclidean distance to all other variants to be infinite, essentially using a single variant test for this variant.

Finally, although we focus on protein structure 3D structure here, we note that the POINT framework may also be extended to incorporate other types of structural information, such as chromosome looping structure from 3C or Hi-C technology, via specifying pairwise distance matrix or correlation matrix among loci.

## Acknowledgments

The authors deeply thank Dr. Peter Vollenweider and Dr. Gerard Waeber, PIs of the CoLaus study, and Dr. Meg Ehm and Dr. Matthew Nelson, collaborators at GlaxoSmithKline, for providing the CoLaus sequence data.

## Supporting information

**S1 Fig. *PLA2G7* variant D69 borrowing map.** Variant-borrowing map defining the amount of borrowing from neighboring variants for *PLA2G7* variant D69 for different values of *c*, with darker color representing higher levels of contribution via the variant correlation matrix *R*.

**S2 Fig. *PLA2G7* variant R82 borrowing map.** Variant-borrowing map defining the amount of borrowing from neighboring variants for *PLA2G7* variant R82 for different values of *c*, with darker color representing higher levels of contribution via the variant correlation matrix *R*.

**S3 Fig. *PLA2G7* variant F110 borrowing map.** Variant-borrowing map defining the amount of borrowing from neighboring variants for *PLA2G7* variant F110 for different values of *c*, with darker color representing higher levels of contribution via the variant correlation matrix *R*.

**S4 Fig. *PLA2G7* variant D181 borrowing map.** Variant-borrowing map defining the amount of borrowing from neighboring variants for *PLA2G7* variant D181 for different values of *c*, with darker color representing higher levels of contribution via the variant correlation matrix *R*.

**S5 Fig. *PLA2G7* variant T187 borrowing map.** Variant-borrowing map defining the amount of borrowing from neighboring variants for *PLA2G7* variant T187 for different values of *c*, with darker color representing higher levels of contribution via the variant correlation matrix *R*.

**S6 Fig. *PLA2G7* variant K191 borrowing map.** Variant-borrowing map defining the amount of borrowing from neighboring variants for *PLA2G7* variant K191 for different values of *c*, with darker color representing higher levels of contribution via the variant correlation matrix *R*.

**S7 Fig. *PLA2G7* variant D200 borrowing map.** Variant-borrowing map defining the amount of borrowing from neighboring variants for *PLA2G7* variant D200 for different values of *c*, with darker color representing higher levels of contribution via the variant correlation matrix *R*.

**S8 Fig. *PLA2G7* variant S273 borrowing map.** Variant-borrowing map defining the amount of borrowing from neighboring variants for *PLA2G7* variant S273 for different values of *c*, with darker color representing higher levels of contribution via the variant correlation matrix *R*.

**S9 Fig. *PLA2G7* variant L283 borrowing map.** Variant-borrowing map defining the amount of borrowing from neighboring variants for *PLA2G7* variant L283 for different values of *c*, with darker color representing higher levels of contribution via the variant correlation matrix *R*.

**S10 Fig. *PLA2G7* variant G303 borrowing map.** Variant-borrowing map defining the amount of borrowing from neighboring variants for *PLA2G7* variant G303 for different values of *c*, with darker color representing higher levels of contribution via the variant correlation matrix *R*.

**S11 Fig. *PLA2G7* variant A326 borrowing map.** Variant-borrowing map defining the amount of borrowing from neighboring variants for *PLA2G7* variant A326 for different values of *c*, with darker color representing higher levels of contribution via the variant correlation matrix *R*.

**S12 Fig. *PLA2G7* variant M331 borrowing map.** Variant-borrowing map defining the amount of borrowing from neighboring variants for *PLA2G7* variant M331 for different values of *c*, with darker color representing higher levels of contribution via the variant correlation matrix *R*.

**S13 Fig. Quantile-quantile plots of p-values for *PLA2G7* simulation study under the null hypothesis of no causal variants using different methods.**

SVT: single variant test; POINT-Burden: POINT test using local burden kernel; POINT-Linear: POINT test using local linear kernel; SCAN: scan statistic method (from the p-values of the best window). The top panel is for continuous traits and the bottom panel is for binary traits.

**S14 Fig. Empirical receiver operating characteristic (ROC) curves for quantitative analysis under different scenarios of causal variants.** The simulation scenarios are listed in Tables 2 and 3. The Y-axis is the true positive rate (i.e., sensitivity) and the X-axis is the false positive rate (i.e., 1-specificity). Red dotted line: single variant test (SVT); blue dashed line: POINT test using local burden kernel; green dash-dot line: POINT test using local linear kernel.

**S15 Fig. Empirical receiver operating characteristic (ROC) curves for binary analysis under different scenarios of causal variants.** The simulation scenarios are listed in Tables 2 and 3. The Y-axis is the true positive rate (i.e., sensitivity) and the X-axis is the false positive rate (i.e., 1-specificity). Red dotted line: single variant test; blue dashed line: POINT test using local burden kernel (SVT); green dash-dot line: POINT test using local linear kernel; purple short-dash line: scan statistic method (SCAN).

**S16 Fig. *PCSK9* variant-borrowing map**. Variant-borrowing map showing the amount of borrowing from neighboring variants for the chosen best *c* value for the local kernel test of association between the rare variants in *PCSK9* and LDL (for conciseness, variants with *c* = 0 are not shown).

**S1 Table. *PLA2G7* rare variant summary information.** Minor allele frequency and protein coordinate information for the rare variants in *PLA2G7*. The 3D coordinates are obtained from PDB entry 3F96.

**S2 Table. *PCSK9* rare variant summary information.** Minor allele frequency and protein coordinate information for the rare variants in *PCSK9*. The 3D coordinates are obtained from PDB entry 4K8R.

**S1 Appendix. Resampling Approach to Obtain the P-value of the Localized Test for Variant** *m*.

